# Sensitivity of novel SARS-CoV-2 Omicron subvariants, BA.2.11, BA.2.12.1, BA.4 and BA.5 to therapeutic monoclonal antibodies

**DOI:** 10.1101/2022.05.03.490409

**Authors:** Daichi Yamasoba, Yusuke Kosugi, Izumi Kimura, Shigeru Fujita, Keiya Uriu, Jumpei Ito, Kei Sato, The Genotype to Phenotype Japan (G2P-Japan) Consortium

**Affiliations:** The University of Tokyo, Tokyo, Japan; Kobe University, Hyogo, Japan

**Author notes:** Correspondence (Kei Sato). Dr. Yamasoba, Mr. Kosugi and Dr. Kimura contributed equally to this letter.

## Abstract

As of May 2022, Omicron BA.2 variant is the most dominant variant in the world. Thereafter, Omicron subvariants have emerged and some of them began outcompeting BA.2 in multiple countries. For instance, Omicron BA.2.11, BA.2.12.1 and BA.4/5 subvariants are becoming dominant in France, the USA and South Africa, respectively. In this study, we evaluated the sensitivity of these new Omicron subvariants (BA.2.11, BA.2.12.1 and BA.4/5) to eight therapeutic monoclonal antibodies (bamlanivimab, bebtelovimab, casirivimab, cilgavimab, etesevimab, imdevimab, sotrovimab and tixagevimab). Notably, we showed that although cilgavimab is antiviral against BA.2, BA.4/5 exhibits higher resistance to this antibody compared to BA.2. Since mutations are accumulated in the spike proteins of newly emerging SARS-CoV-2 variants, we suggest the importance of rapid evaluation of the efficiency of therapeutic monoclonal antibodies against novel SARS-CoV-2 variants.

## Text

During the current pandemic, severe acute respiratory syndrome coronavirus 2 (SARS-CoV-2) has considerably diversified. The Omicron variant was identified at the end of November 2021 and rapidly spread worldwide. As of May 2022, Omicron BA.2 variant is the most dominant variant in the world. Thereafter, Omicron subvariants have emerged and some of them began outcompeting BA.2 in multiple countries. For instance, Omicron BA.2.11, BA.2.12.1 and BA.4/5 subvariants are becoming dominant in France, the USA and South Africa, respectively (**Figure 1A**).

**Figure 1.**
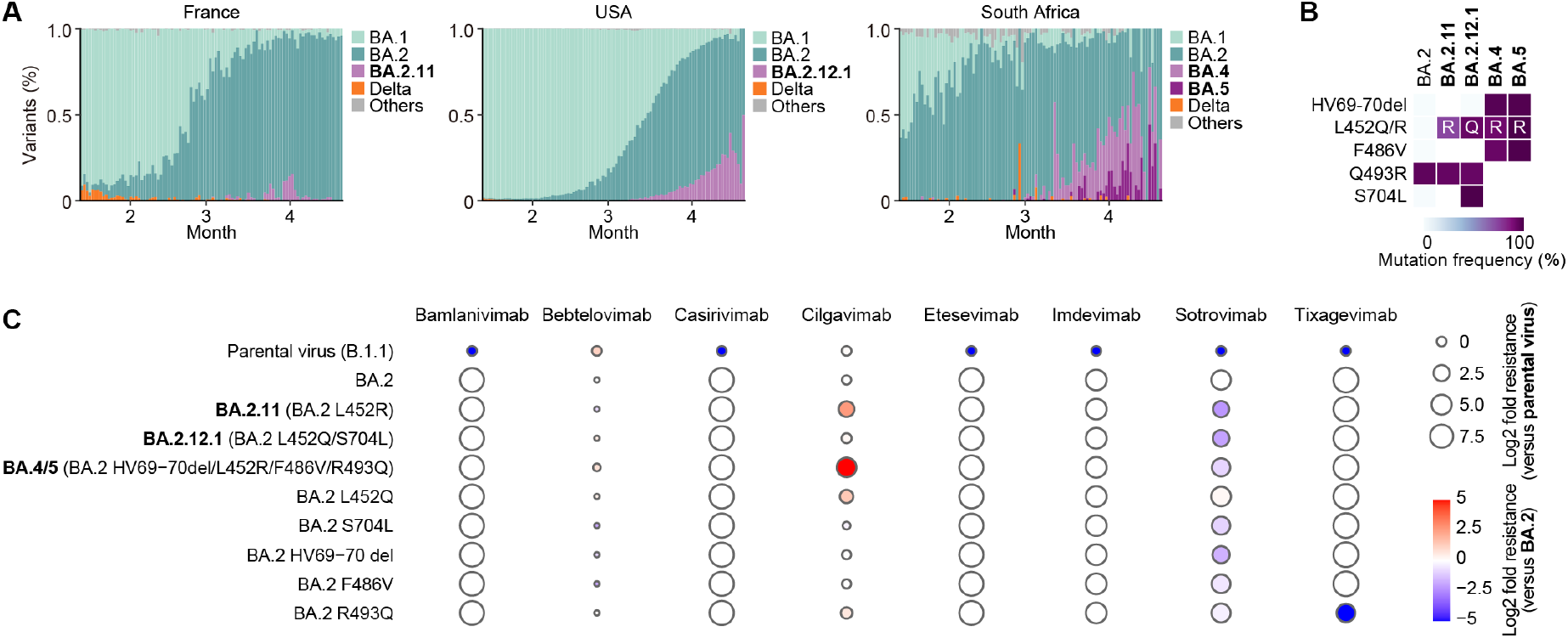
Sensitivity of new Omicron subvariants to eight therapeutic monoclonal antibodies. (**A**) Epidemics of SARS-CoV-2 lineages in in France, the USA and South Africa. The data from January 13, 2022 to April 22, 2022 (100 days) for France, the USA and South Africa were analyzed. In this figure, the SARS-CoV-2 lineages other than Delta and Omicron are shown as “others”. (**B**) Amino acid substitutions in S. Heatmap shows the frequency of amino acid substitutions in BA.2.11, BA.2.12.1, BA.4, and BA.5 compared to BA.2. Substitutions detected in >50% of sequences of any lineage are shown. (**C**) Virus neutralization assays. A neutralization assay was performed using pseudoviruses harboring the SARS-CoV-2 S proteins of Omicron subvariants [BA.2.11 (BA.2 S:L452R), BA.2.12.1 (BA.2 S:L452Q/S704L) and BA.4/5 (BA.2 S: HV69-70del/L452R/F486V/R493Q)], their derivatives (the BA.2 S bearing L452Q, S704L, HV69-70del, F486V or R493Q, respectively) or the D614G-harboring B.1.1 lineage virus (parental virus). Eight therapeutic monoclonal antibodies (bamlanivimab, bebtelovimab, casirivimab, cilgavimab, etesevimab, imdevimab, sotrovimab and tixagevimab) were tested. The assay of each antibody was performed in sextuplicate at each concentration to determine the 50% neutralization concentration. The log2 fold changes of resistance versus the parental virus (circle size) or BA.2 (color) are respectively shown. Representative neutralization curves are shown in **Figure S1** in the Supplementary Appendix.

Newly emerging SARS-CoV-2 variants need to be carefully monitored for a potential increase in transmission rate, pathogenicity and/or resistance to immune responses. The resistance of variants to vaccines and therapeutic antibodies can be attributed to a variety of mutations in the viral spike (S) protein. Although the S proteins of new Omicron subvariants (BA.2.11, BA.2.12.1 and BA.4/5) are based on the BA.2 S, the majority of them additionally bear the following substitutions in the S: BA.2.11, L452R; BA.2.12.1, L452Q and S704L; and BA.4/5, L452R, HV69-70del, F486V and R493Q (**Figure 1B**). In particular, the L452R and L452Q substitutions were detected in Delta and Lambda variants, and we demonstrated that the L452R/Q substitution affects the sensitivity to vaccine-induced neutralizing antibodies.^1,2^ Therefore, it is reasonable to assume that these new Omicron subvariants reduces sensitivity towards therapeutic monoclonal antibodies. To address this possibility, we generated pseudoviruses harboring the S proteins of these Omicron subvariants and derivatives and prepared eight therapeutic monoclonal antibodies. Consistent with previous studies,^3-5^ bamlanivimab, casirivimab, etesevimab, imdevimab and tixagevimab were not functional against BA.2 (**Figure 1C**). These five antibodies did not work against new Omicron subvariants, while the BA.2 S bearing R493Q substitution was partially sensitive to casirivimab and tixagevimab (**Figure 1C** and **Figure S1**). Interestingly, bebtelovimab was ∼2-fold more effective against BA.2 and all Omicron subvariants tested than the parental virus (**Figure 1C**). Although sotrovimab was ∼20-fold less antiviral against BA.2 than the parental virus, the Omicron subvariants bearing L452R substitution including BA.2.11 and BA.4/5 were more sensitive to sotrovimab than BA.2 (**Figure 1C**). Cilgavimab was also antiviral against BA.2, while the L452R/Q substitution rendered ∼2-5-fold resistance to this antibody. Notably, BA.4/5 exhibited ∼30-fold more resistance to cilgavimab compared to BA.2 (**Figure 1C**).

Since mutations are accumulated in the S proteins of newly emerging SARS-CoV-2 variants, we suggest the importance of rapid evaluation of the efficiency of therapeutic monoclonal antibodies against novel SARS-CoV-2 variants.

## Grants

Supported in part by AMED Research Program on Emerging and Re-emerging Infectious Diseases 20fk0108146 (to Kei Sato), 20fk0108270 (to Kei Sato), 20fk0108413 (to Kei Sato) and 20fk0108451 (to G2P-Japan Consortium and Kei Sato); AMED Research Program on HIV/AIDS 21fk0410039 (to Kei Sato); JST SICORP (e-ASIA) JPMJSC20U1 (to Kei Sato); JST SICORP JPMJSC21U5 (to Kei Sato), JST CREST JPMJCR20H4 (to Kei Sato); JSPS KAKENHI Grants-in-Aid for Scientific Research B 18H02662 (to Kei Sato) and 21H02737 (to Kei Sato); JSPS Fund for the Promotion of Joint International Research (Fostering Joint International Research) 18KK0447 (to Kei Sato); JSPS Core-to-Core Program JPJSCCA20190008 (A. Advanced Research Networks) (to Kei Sato); JSPS Research Fellow DC1 19J20488 (to Izumi Kimura) and 22J11578 (to Keiya Uriu); The Tokyo Biochemical Research Foundation (to Kei Sato); and Joint Usage/Research Center program of Institute for Frontier Life and Medical Sciences, Kyoto University (to Kei Sato).

## Notes

**Conflict of interest**: The authors declare that no competing interests exist.

### Competing Interest Statement

The authors have declared no competing interest.

