## Supplementary information for "Sensitivity of novel SARS-CoV-2 Omicron subvariants, BA.2.11, BA.2.12.1, BA.4 and BA.5 to therapeutic monoclonal antibodies"

**Supplementary Appendix**

**Table of Contents**

**Contents** Page

| | |
|---|---|
| **Materials and Methods** | 2-3 |
| Epidemics and mutation analysis | |
| Cell culture | |
| Plasmid construction | |
| Preparation of monoclonal antibodies | |
| Neutralization assay | |
| **Figure S1.** Representative neutralization curves. | 4 |
| **Table S1.** Primers used for the construction of SARS-CoV-2 S expression plasmids. | 5 |
| **Consortia** | 6 |
| **Acknowledgments** | 7 |
| **Supplemental References** | 8 |

### Materials and Methods

#### Epidemics and mutation analysis

The genome surveillance data of SARS-CoV-2 was downloaded from the GISAID database (https://www.gisaid.org/) on April 29, 2022. We excluded the data of viral strains with the following features from the analysis: i) a lack of collection date information; ii) sampling in animals other than humans; or iii) sampling by quarantine. The data for South Africa, France, and the USA from January 13, 2022 to April 22, 2022 (100 days) were analyzed. The epidemic dynamics of BA.1, BA.2, Delta, and the Omicron lineages harboring L452R or L452Q mutations in spike (e.g., BA.2.11, BA.2.12.1, BA.4 and BA.5) are shown in **Figure 1A**. In this figure, the SARS-CoV-2 lineages other than above were summarized as "others".

Amino acid differences among the S proteins of BA.2, BA.2.11, BA.2.12.1, BA.4, and BA.5 (**Figure 1B**) were extracted from the GISAID data above as follows: in the SARS-CoV-2 lineages above as well as BA.2, the prevalence of amino acid substitutions in spike compared to hCoV-19/Wuhan/WIV04/2019 (GISAID ID: EPI_ISL_402124) spike were calculated. Subsequently, amino acid substitutions that were detected in any lineages with >50% prevalence were extracted. Finally, amino acid substitutions shared in all lineages above were excluded.

#### Cell culture

HEK293T cells (a human embryonic kidney cell line; ATCC CRL-3216) and HOS-ACE2/TMPRSS2 cells (kindly provided by Dr. Kenzo Tokunaga),[1,2] a derivative of HOS cells (a human osteosarcoma cell line; ATCC CRL-1543) stably expressing human ACE2 and TMPRSS2, were maintained in Dulbecco's modified Eagle's medium (DMEM) (high glucose) (Wako, Cat# 044-29765) containing 10% fetal bovine serum (FBS) (Sigma-Aldrich Cat# 172012-500ML), 100 units penicillin and 100 ug/ml streptomycin (PS) (Sigma-Aldrich, Cat# P4333-100ML).

#### Plasmid construction

To construct the plasmids expressing anti-SARS-CoV-2 monoclonal antibodies (bamlanivimab, bebtelovimab, casirivimab, cilgavimab, etesevimab, imdevimab, sotrovimab and tixagevimab), the sequences of the variable regions of these antibodies were obtained from KEGG Drug Database (https://www.genome.jp/kegg/drug/) and were artificially synthesized by Fasmac. The obtained coding sequences of the variable regions of the heavy and light chains were cloned into the pCAGGS vector containing the sequences of the human immunoglobulin 1 and kappa constant region (kindly provided by Dr. Hisashi Arase). Plasmids expressing the SARS-CoV-2 spike proteins of the parental D614G (B.1.1) and Omicron BA.2 were prepared in our previous studies.[2,3] Plasmids expressing the spike protein of Omicron variants (BA.2.11, BA.2.12.1 and BA.4/5) and their derivatives were generated by site-directed overlap extension PCR using pC-SARS2-S D614G[2] as the template and the primers listed in **Table S1**. The resulting PCR fragment was subcloned into the KpnI-NotI site of the pCAGGS vector[4] using In-Fusion® HD Cloning Kit (Takara, Cat# Z9650N). Nucleotide sequences were determined by DNA sequencing services (Eurofins), and the sequence data were analyzed by Sequencher v5.1 software (Gene Codes Corporation).

**Preparation of monoclonal antibodies**

Eight monoclonal antibodies (bamlanivimab, bebtelovimab, casirivimab, cilgavimab, etesevimab, imdevimab, sotrovimab and tixagevimab) were prepared as previously described.[3,5,6] Briefly, the pCAGGS vectors containing the sequences encoding the immunoglobulin heavy and light chains were cotransfected into HEK293T cells at 1:1 ratio using PEI Max (Polysciences, Cat# 24765-1). The culture medium was refreshed with DMEM (low glucose) (Wako, Cat# 041-29775) containing 10% FBS without PS. At 96 h posttransfection, the culture medium was harvested, and the antibodies were purified using NAb protein A plus spin kit (Thermo Fisher Scientific, Cat# 89948) according to the manufacturer's protocol.

**Neutralization assay**

Pseudoviruses were prepared as previously described.[1,3,7-12] Briefly, lentivirus (HIV-1)-based, luciferase-expressing reporter viruses were pseudotyped with the SARS-CoV-2 spikes. HEK293T cells (1 × $10^6$ cells) were cotransfected with 1 μg psPAX2-IN/HiBiT,[13] 1 μg pWPI-Luc2,[13] and 500 ng plasmids expressing parental S or its derivatives using PEI Max (Polysciences, Cat# 24765-1) according to the manufacturer's protocol. Two days post transfection, the culture supernatants were harvested and centrifuged. The pseudoviruses were stored at –80°C until use.

Neutralization assays were performed as previously described.[3,6,7,9,11,12] Briefly, the SARS-CoV-2 spike pseudoviruses (counting ~20,000 relative light units) were incubated with serially diluted monoclonal antibodies at 37°C for 1 h. Pseudoviruses without monoclonal antibody were included as controls. Then, an 40 μl mixture of pseudovirus and serum was added to HOS-ACE2/TMPRSS2 cells (10,000 cells/50 μl) in a 96-well white plate. Two days post infection, the infected cells were lysed with a Bright-Glo luciferase assay system (Promega, Cat# E2620), and the luminescent signal was measured using a GloMax explorer multimode microplate reader 3500 (Promega). The assay of each monoclonal antibody was performed in triplicate, and the 50% neutralization titer was calculated using Prism 9 (GraphPad Software).

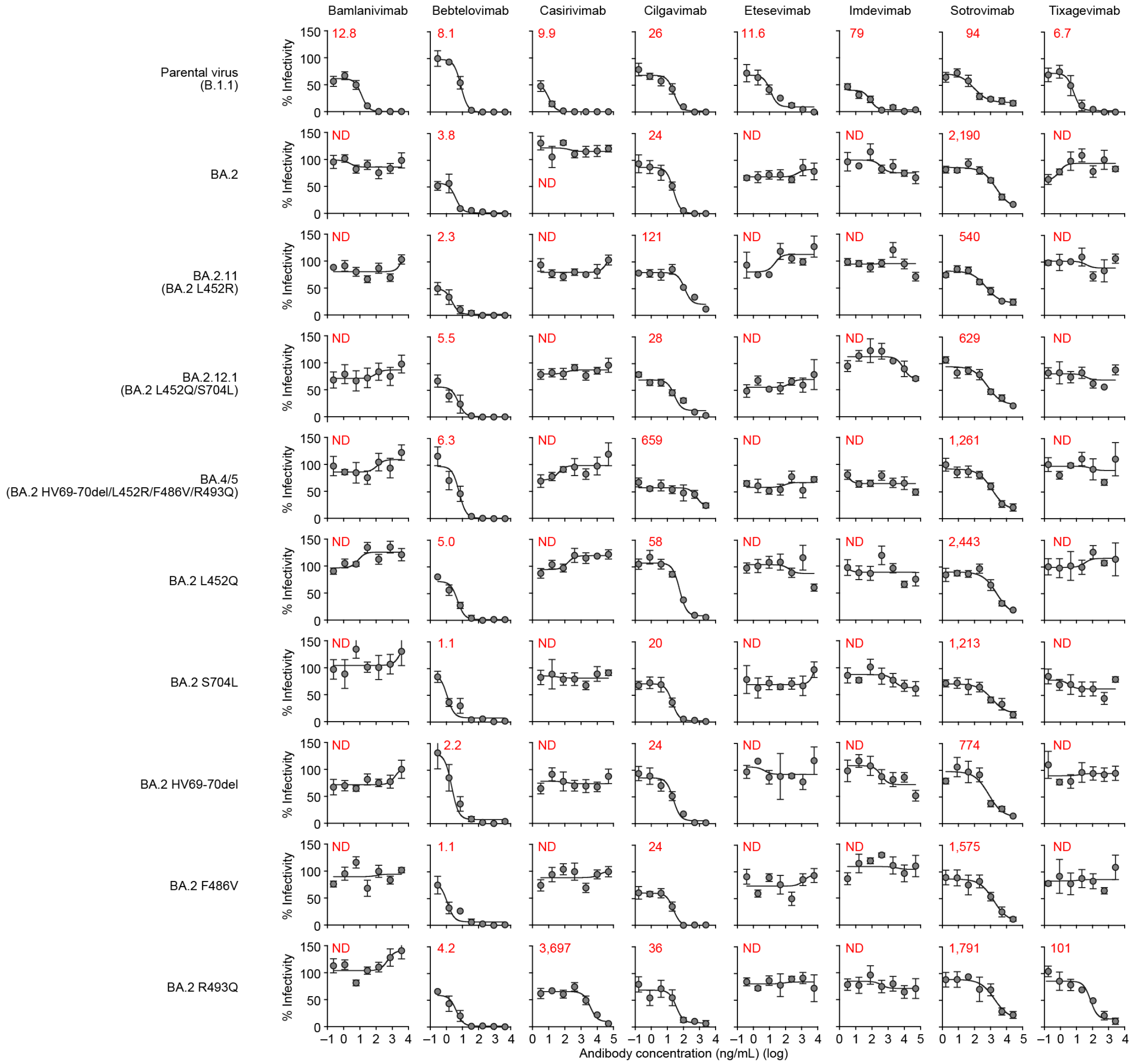


**Figure S1. Representative neutralization curves.** A neutralization assay was performed using pseudoviruses harboring the SARS-CoV-2 spike proteins of Omicron subvariants [BA.2.11 (BA.2 S:L452R), BA.2.12.1 (BA.2 S:L452Q/S704L) and BA.4/5 (BA.2 S: HV69-70del/L452R/F486V/R493Q)], their derivatives (the BA.2 S bearing L452Q, S704L, HV69-70del, F486V or R493Q, respectively) or the D614G-harboring B.1 lineage virus (parental virus). Eight therapeutic monoclonal antibodies (bamlanivimab, bebtelovimab, casirivimab, cilgavimab, etesevimab, imdevimab, sotrovimab and tixagevimab) were tested. The assay of each antibody was performed in sextuplicate at each concentration to determine the 50% neutralization concentration (ng/mL). The presented data are expressed as the average with standard deviation, and representative neutralization curves are shown. The red numbers in the panels indicate the 50% neutralization concentration (ng/mL). ND, not determined. Summarized data are shown in **Figure 1C**.

**Table S1. Primers used for the construction of SARS-CoV-2 S expression plasmids.**

| Primer name | Sequence (5'-to-3') |
|---|---|
| Omicron universal Fw | cactatagggcgaattgggtaccatgtttgtgttcctggt |
| BA2 Rv | agctccaccgcggtggcggccgctcaggtgtagtgcagtttca |
| pC-S_BA2_L452M-F | caactacaactacatgtacagactgttca |
| pC-S_BA2_L452M-R | tgaacagtctgtacatgtagttgtagttg |
| pC-S_BA2_L452Q-F | caactacaactaccagtacagactgttca |
| pC-S_BA2_L452Q-R | tgaacagtctgtactggtagttgtagttg |
| pC-S_BA2_L452Q_setSL-F | gggagcagagaacctggtggcttacagca |
| pC-S_BA2_L452Q_setSL-R | tgctgtaagccaccaggttctctgctccc |
| OptS L452R F | caactacaactaccgttacagactgttcagg |
| OptS L452R R | cctgaacagtctgtaacggtagttgtagttg |
| pC-S_BA2_6970del_F | tggttccatgccatctctggcaccaatggc |
| pC-S_BA2_6970del_R | gccattggtgccagagatggcatggaacca |
| pC-S_BA2_F486V-F | tggagtggccggcgtgaactgttactttc |
| pC-S_BA2_F486V-R | gaaagtaacagttcacgccggccactcca |
| pC-S_BA2_R493Q-F | ttactttccactccaatcctatggcttca |
| pC-S_BA2_R493Q-R | tgaagccataggattggagtggaaagtaa |
| pC-S_BA2_F486V_R493Q-F | tggccggcgtgaactgttactttccactccaatcctatgg |
| pC-S_BA2_F486V_R493Q-R | ccataggattggagtggaaagtaacagttcacgccggcca |

### Consortia

#### The Genotype to Phenotype Japan (G2P-Japan) Consortium

**The Institute of Medical Science, The University of Tokyo, Japan**
Mai Suganami, Mika Chiba, Ryo Yoshimura, Naoko Misawa

**Hokkaido University, Japan**
Takasuke Fukuhara, Keita Matsuno, Shinya Tanaka, Rigel Suzuki, Tomokazu Tamura, Yuhei Morioka, Kana Tsushima, Haruko Kubo, Mai Kishimoto, Hirofumi Sawa, Naganori Nao, Masumi Tsuda, Lei Wang, Yoshitaka Oda

**Tokyo Metropolitan Institute of Public Health, Japan**
Kenji Sadamasu, Kazuhisa Yoshimura, Hiroyuki Asakura, Isao Yoshida, Mami Nagashima

**Tokai University, Japan**
So Nakagawa, Jiaqi Wu, Miyoko Takahashi

**Kyoto University, Japan**
Kotaro Shirakawa, Akifumi Takaori-Kondo, Yasuhiro Kazuma, Ryosuke Nomura, Yoshihito Horisawa

**Hiroshima University, Japan**
Takashi Irie, Ryoko Kawabata

**Kumamoto University, Japan**
Terumasa Ikeda, Chihiro Motozono, Mako Toyoda, Takamasa Ueno, Hesham Nasser, Ryo Shimizu, Kazuko Kitazato, Haruyo Hasebe, Toong Seng Tan

**University of Miyazaki, Japan**
Akatsuki Saito, Shuya Mitoma, Erika P Butlertanaka, Yuri L Tanaka

**Acknowledgments**

We would like to thank all members of The Genotype to Phenotype Japan (G2P-Japan) Consortium. We thank Dr. Kenzo Tokunaga (National Institute of Infectious Diseases, Japan) and Dr. Hisashi Arase (Osaka University, Japan) for sharing materials. We gratefully acknowledge the numerous laboratories worldwide that have provided sequence data and metadata to GISAID. A full list of originating and submitting laboratories for the sequences used in our analysis can be found at https://www.gisaid.org using the EPI-SET-ID: EPI_SET_20220503qc.
